# Structure-function conservation between the methyltransferases SETD3 and SETD6

**DOI:** 10.1101/2022.03.31.486554

**Authors:** Lee Elisha, Elina Abaev-Schneiderman, Ofir Cohn, Guy Shapira, Noam Shomron, Michal Feldman, Dan Levy

## Abstract

Among the protein lysine methyltransferases family members, it appears that SETD6 is highly similar and closely related to SETD3. The two methyltransferases show high similarity in their structure, which raised the hypothesis that they share cellular functions. Using a proteomic screen, we identified 52 shared interacting-proteins. Gene Ontology (GO) analysis of the shared proteins revealed significant enrichment of proteins involved in transcription. Our RNA-seq data of SETD6 KO and SETD3 KO HeLa cells identified ∼100 up-regulated and down-regulated shared genes. We have also identified a substantial number of genes that changed dramatically in the double KO cells but did not significantly change in the single KO cells. GO analysis of these genes revealed enrichment of apoptotic genes. Accordingly, we show that the double KO cells displayed high apoptotic levels, suggesting that SETD6 and SETD3 inhibit apoptosis. Collectively, our data strongly suggest a functional link between SETD6 and SETD3 in the regulation of apoptosis.

## Introduction

Methylation of lysine (K) residues is mediated by a group of enzymes known as protein methyltransferases (PKMT). There are over 60 members in this enzyme family, the majority of them belong to the SET domain-containing group, which is the domain responsible for the enzymatic activity [1, 2]. The second group includes only a few characterized human PKMTs which belong to the seven-β-strand (7βS) methyltransferase family [3]. The ε -amino group of lysine can be mono-di- or tri-methylated in an S-adenosyl methionine-dependent manner. S-Adenosyl methionine (SAM) is the major methyl donor in the cell, and it is the second most frequently used substrate, following ATP [4]. The large number of methyltransferases and the abundance of SAM in the cell point to the importance of the methylation reaction in various cellular processes [5].

SET domain-containing protein 6-SETD6 is a member of the lysine methyltransferase family. It is a mono-methyltransferase which contains a catalytic SET domain and a Rubisco substrate-binding domain. SETD6 was identified as a regulator of inflammation through the methylation of NF-kB/RelA protein [6]. Later studies revealed its role in a variety of cellular processes, mostly through the regulation of transcription. For example, it was found to methylate the histone H2A variant H2AZ to maintain embryonic stem-cell renewal [7, 8]. It was also found to regulate the NRF2-mediated oxidative stress response, the WNT/β-catenin target genes and the nuclear hormone receptor signaling [7, 9, 10]. Recently, SETD6 was found to regulate the protein-translation process through the methylation of BRD4 [11]. In addition, SETD6 was found to regulate mitotic progression in a non-transcriptional manner through the methylation of PLK1 (Polo-like kinase 1), a master cell-cycle regulator [12]

SETD3 is also a member of the methyltransferase family. Among the members of the protein lysine methyltransferases family, SETD3 is highly similar and closely related to SETD6 [13]. Similar to SETD6, SETD3 contains a catalytic SET domain and a Rubisco substrate-binding domain. SETD3 was identified in 2011 and characterized as a histone H3 methyltransferase in Zebrafish and mice [14]. It has high expression levels in muscle tissue, where it promotes myocyte differentiation [15]. Under hypoxic conditions, SETD3 methylates the transcription factor FOXM1 to repress VEGF levels [16]. SETD3 was also found as a positive regulator of DNA-damage-induced apoptosis by regulating p53 target genes [17]. Other studies have suggested SETD3 as a prognostic biomarker for Hepatocellular carcinoma cell (HCC), breast cancer, renal cell tumors, early-stage ovarian carcinoma, and colon cancer [17-21]. Recently, several studies reported SETD3 as the first mammalian histidine methyltransferase, which catalyzes His73 methylation of actin [22, 23].

The functional cellular redundancy in the lysine methyltransferase family was studied in depth mainly in the context of histone proteins. For example, Suv39H1 and Suv39H2 were shown to display redundant roles: disruption of one gene did not affect embryonic development, while disruption of both severely impaired viability [24]. GLP and G9a di-methylate H3K9. However, this activity is not redundant as both are required for mammalian development [25], and GLP could not compensate for G9a loss in human rectal cancer cells [26]. In addition, all 6 members of the MLL family methylate H3K4. However, they display different activities in a tissue and developmental stages dependent manners [27].

The two closely related methyltransferases SETD3 and SETD6 show high similarity in their structure, which raised the hypothesis that they share common functions in cells. In this study, we show that both enzymes have common interacting proteins and target genes. Specifically, we found that cells depleting both enzymes present a high level of cell-death-related genes and are more sensitive to DNA-damage-induced apoptosis. Cells depleting only one of the enzymes did not exhibit this effect, suggesting that they both negatively regulate apoptosis and can compensate for the depletion of each other. Altogether, we present here a functional link between SETD3 and SETD6 in the regulation of apoptosis.

## Methods

### Cell Lines, Plasmids, Treatments

Human cervical cancer (HeLa) cells were maintained in Dulbecco’s modified Eagle’s medium (Sigma, D5671) with 10% fetal bovine serum (FBS) (Gibco), penicillin-streptomycin (Sigma, P0781), 2 mg/ml L-glutamine (Sigma, G7513) and non-essential amino acids (Sigma, M7145), at 37°C in a humidified incubator with 5% CO_2_. HeLa CRISPR/Cas9 SETD3 KO and SETD6 KO cells were generated as previously described [12, 16], HeLa CRISPR-Cas9 double KO cells were generated from SETD6 KO cells using SETD3 KO gRNAs. For control cells, lentiCRISPR plasmid with no gRNAs was used. Plasmids used for expression and purification of recombinant proteins were His Sumo-SETD3 and His SETD6, previously described in [11, 16].

For Doxorubicin treatment, cells were seeded in 12-well plates and 2μM Doxorubicin was added for 24h; DMSO was used as vehicle.

### Plasmids and Recombinant Proteins expression and purification

For recombinant purification SETD6 and SETD3 cDNA were subcloned into pET-Duet plasmids. *Escherichia coli* Rosetta strain (Novagen) transformed with a plasmid expressing His-tagged SETD3 or SETD6 proteins were grown in LB medium. Bacteria were harvested by centrifugation after IPTG induction and lysed by sonication on ice (25% amplitude, 1 min total, 10/5 sec ON/OFF). His-tagged proteins were purified using Ni-NTA beads (Pierce) on a HisTrap column (GE) with the ÄKTA gel filtration system. Proteins were eluted by 0.5 M imidazole followed by dialysis to 10% glycerol in phosphate-buffered saline (PBS).

### Antibodies, Western Blot Analysis

Primary antibodies used were: anti-Actin (Abcam, ab3280), anti-SETD6 (Genetex, GTX629891), Anti-SETD3 (ab176582; Abcam), HRP-conjugated secondary antibodies, goat anti-rabbit, goat anti-mouse, (were purchased from Jackson ImmunoResearch (111-035-144, 115-035-062 respectively). Fluorescently labeled secondary antibodies used: Alexa 647 anti-rabbit and anti-mouse (Invitrogen, A-21443). For Western blot analysis, cells were homogenized and lysed in RIPA buffer (50 mM Tris-HCl pH 8, 150 mM NaCl, 1% Nonidet P-40, 0.5% sodium deoxycholate, 0.1% SDS, 1 mM DTT, and 1:100 protease inhibitor mixture (Sigma)). Samples were resolved on SDS-PAGE, followed by Western blot analysis.

### RNA extraction and quantitative RT-PCR

Total RNA was extracted using the NucleoSpin RNA Kit (Macherey-Nagel). 200 ng of the extracted RNA was reverse-transcribed to cDNA using the iScript cDNA Synthesis Kit (Bio-Rad) according to the manufacturer’s instructions. Real-time qPCR was performed using the UPL probe library system in a LightCycler 480 System (Roche). The real-time qPCR primers were designed using the universal probe library assay design center (Roche). All samples were amplified in triplicates in a 384-well plate using the following cycling conditions: 10 min at 95°C, 45 cycles of 10 sec at 95°C, 30 sec at 60°C and 1 sec at 72°C, followed by 30 sec at 40°C. Gene expression levels were normalized to GAPDH and controls of the experiment.

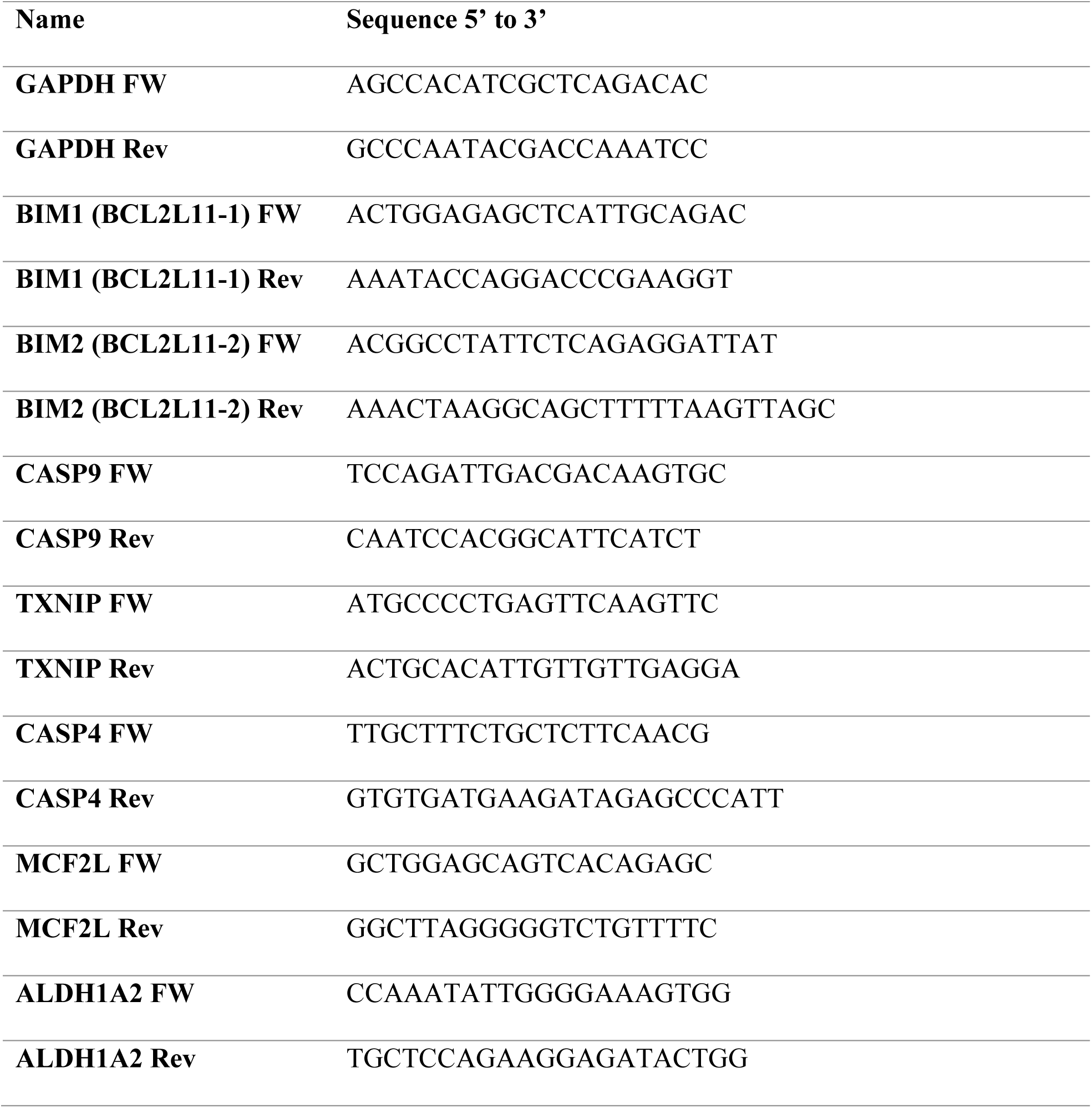

### ProtoArray

Human protein arrays (Version 5.0; ProtoArray) were stored at −20 °C until use. Arrays were blocked with blocking buffer (5 mM MgCl_2_, 0.5 mM DTT, 0.05% Triton X-100, 5% glycerol, 1% BSA, 10% PBS) at room temperature for 1 h. Arrays were then washed with probing buffer (0.1% Tween 20, 1% BSA, 10% PBS) and incubated for 1.5 h in a hybridization chamber (Agilent, Santa Clara, CA) in a reaction mixture containing 80 μg of purified His-SETD6 in probing buffer in a total reaction volume of 950 μl. Arrays were washed four times with probing buffer while shaking at 50 rpm for 5 min at room temperature. Arrays were then incubated with SETD6 antibody for 1 hour at room temperature while shaking at 50 rpm. Arrays were washed four times with probing buffer while shaking at 50 rpm for 5 minutes at room temperature, followed by incubation for 1 h with Alexa Fluor 647 chicken anti-mouse IgG (Invitrogen) diluted in probing buffer while shaking at 50 rpm. The arrays were washed four times as described, followed by one wash with PBSx1 and then with DDW. The arrays were then dried in a centrifuge at 300 rcf for 1 min. Arrays were scanned (Axon GenePix 4000B; Molecular Devices Inc. Sunnyvale, CA, USA) and data were analyzed for each block using software alignment (Genepix Pro 7 software; Molecular Devices, Sunnyvale, CA) and gene array list (GAL) files supplied by the protein array manufacturer (Invitrogen). ProtoArray experiments for His-SETD3 were performed as previously described [28]. For negative control, the arrays were probed with BSA. The signals were analyzed using the ProtoArray Prospector software V5.2 (Invitrogen). Proteins were defined as positive hits when the Z-score was >3.

### Apoptosis Assay by Flow Cytometry (FACS)

Doxorubicin treated cells were collected by trypsin and stained with Annexin V-FITC (1:200) and 4′,6-Diamidine-2′-phenylindole dihydrochloride (DAPI) (1:400) in 80μl binding buffer (100 mM HEPES, 140 mM NaCl, and 25 mM CaCl_2_, pH 7.4) using a MEBCYTO Apoptosis kit (MBL, Nagoya). After 15 min incubation at room temperature in the dark, 120μl binding buffer was added and samples were analyzed by flow cytometry (Guava® easyCyte flow cytometer). Apoptosis was defined as the total percentage of cells positive for both DAPI and Annexin V-FITC.

### Cell Cycle Assay by FACS

For double thymidine block, HeLa cells were plated at 30% confluence, then supplemented with 2mM thymidine (Sigma) for 17 h. Cells were then washed once with PBS, and fresh media was added to the cells for 9 h. For the second block, cells were then supplemented for an additional 15 h with 2mM Thymidine, washed once with PBS, and fresh media was added. Following thymidine release, cells were harvested after 0 and 8h and fixed with 5ml 70% ethanol at −20^0^C (drop-wise under vortex) and incubated at −20^0^C O.N. Cells were than washed with cold PBS and incubated for 1h on ice. Cells were then treated with 200μl staining solution: 50 μg/ml RNAse, 25 μg/ml Propidium-iodide (PI) in PBS 0.1% triton for 20 min at RT in the dark and then analyzed by FACS.

### Cell Proliferation Assay

For the cell proliferation assay, 5×10^4^ cells were seeded on 24-well plates. Cell number was monitored and calculated by a Lionheart™ FX Automated Microscope (4x) every 2 hours. Cell proliferation was calculated for three experimental replicates.

### RNA-Seq and Data Processing

Total RNA was extracted from HeLa cells using the NucleoSpin RNA Kit (Macherey-Nagel). Samples were prepared in duplicates. RNA-seq libraries were prepared at the Crown Genomics institute of the Nancy and Stephen Grand Israel National Center for Personalized Medicine, Weizmann Institute of Science. Libraries were prepared using the INCPM-mRNA-seq protocol. Briefly, the polyA fraction (mRNA) was purified from 500 ng of total input RNA followed by fragmentation and the generation of double-stranded cDNA. Afterwards, a Agencourt Ampure XP beads cleanup (Beckman Coulter), end repair, A base addition, adapter ligation and PCR amplification steps were performed. Libraries were quantified by Qubit (Thermo fisher scientific) and TapeStation (Agilent). Sequencing was done on a Hiseq instrument (Illumina) using two lanes of an SR60_V4 kit, allocating 20M reads per sample (single read sequencing). Initial analysis of the raw sequence reads was carried out using the NeatSeq-Flow platform [29]. The sequences were quality trimmed and filtered using Trim Galore (v0.4.5) and cutadapt (v1.15). Alignment of the reads to the human genome (GRCh38) was done using STAR (v2.5.2a) [30]. The number of reads per gene per sample was counted using RSEM (v1.2.28) [31]. Quality assessment (QA) of the process was carried out using FASTQC (v0.11.8) and MultiQC (v1.0.dev0) [32]. After trimming each sample had an average of 25.8M reads with an average sequence length of 60bp. Statistical testing for identification of differentially expressed genes, gene annotation, clustering and enrichment analysis were performed with the DESeq2 module within the NeatSeq-Flow platform [29]. Gene annotation was done using the “AnnotationHub” R package (snapshot: 2020-04-27). The statistical analysis was done using the DESeq2 [33] R package. For comparison (Contrast) between each of the treatment groups (SETD3 KO, SETD6 KO and DBL KO) and the control (CT), the statistical model considered one factor: the treatment group. The analysis produced p-value, FDR-adjusted p-value and fold of change (FC) per gene for each contrast. Genes with FDR adjusted p-value < 0.05 were considered Differentially Expressed (DE). The RNA-seq data described in this study are deposited to the Gene Expression Omnibus (GEO) repository. The accession number for the RNA-seq data reported in this paper is GEO: GSE199250.

### Bioinformatic Analysis

Shared proteins from the ProtoArray and differentially expressed genes identified in the RNAseq were analyzed by the DAVID tool for biological processes. Structural alignment of SETD3 and SETD6 (PDB ID: 3SMT and 3QXY respectively) was performed using the PyMol software. Sequence alignment of SETD3 and SETD6 amino acid sequences was performed using UniProt alignment tool for sequence similarity (Accession #: Q86TU7 and Q8TBK2, respectively).

## Results

### SETD3 and SETD6 Share High Sequence and Structure Similarity

Previous phylogenetic analysis has revealed that among the protein lysine methyltransferases family, SETD6 is highly similar and closely related to SETD3 [13, 34]. To further characterize the relationship between the two methyltransferases, we performed structure and amino acids sequence alignment. The two enzymes show high resemblance in their fold (Figure 1A) and have 56%-57% similarity in the amino acid sequences of their SET and Rubisco-substrate binding domain, respectively (Figure 1B). These findings suggest that SETD3 and SETD6 may share common biochemical features.

**Figure 1:**
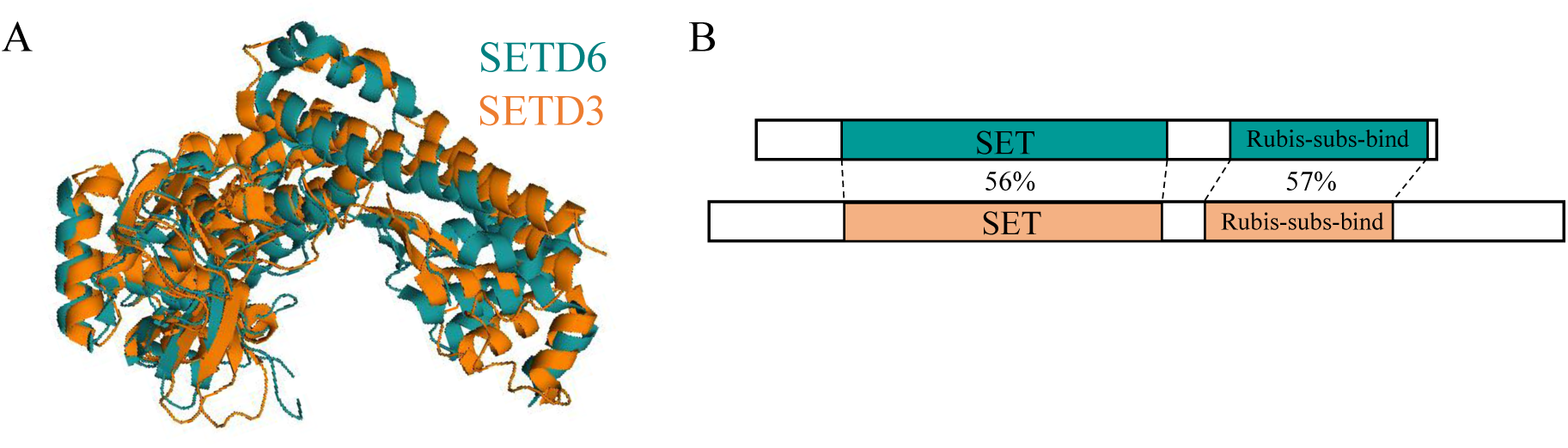
Structure and sequence similarity of SETD6 and SETD3. (**A**) Structure alignment of SETD3 and SETD6 (PDB ID: 3SMT and 3QXY respectively). The alignment performed using the PyMOL software. (**B**) Schematic representation of SETD3 and SETD6 amino-acid sequence alignment with the indicated domains and % similarity. Alignment was performed using the UniProt Align tool.

### SETD3 and SETD6 Share Common Interacting-Proteins

The high biochemical similarity and conservation of SETD3 and SETD6 raised the hypothesis that they share common interacting proteins. To address this hypothesis, we have performed a proteomic screen using the ProtoArray system (ProtoArray®; Invitrogen Corp., Carlsbad, CA, USA). This array contains more than 9,500 highly purified recombinant human proteins, expressed in insect cells as N-terminal glutathione S-transferase (GST) fusion proteins, which are immobilized in duplicates at spatially addressable positions on nitrocellulose-coated glass microscope slides [6]. The arrays were probed with either recombinant His-SETD3 or His-SETD6, followed by anti-His antibody and 647 Alexa fluor-conjugated secondary antibodies (Figure 2A).

**Figure 2.**
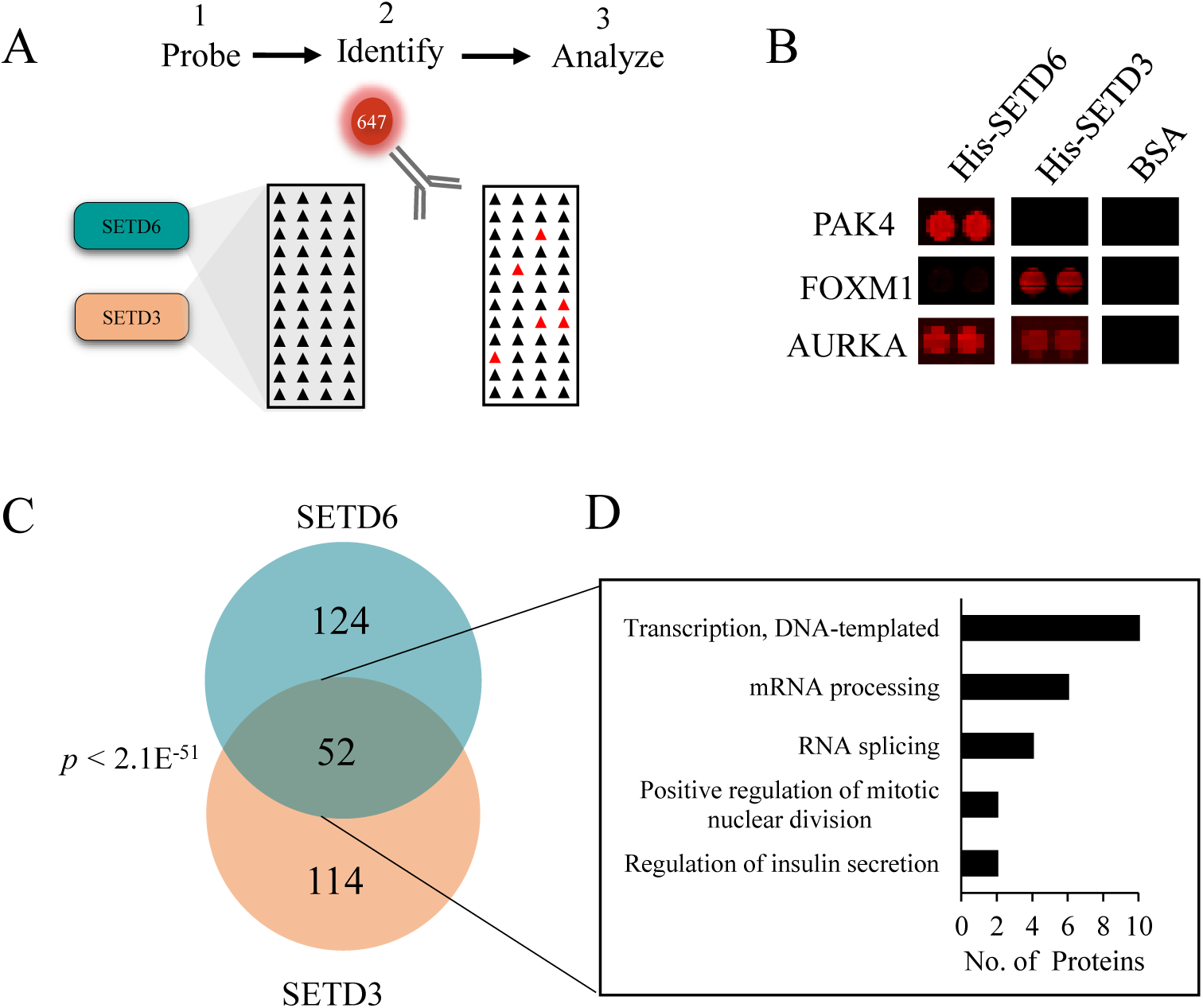
SETD3 and SETD6 share have common interacting proteins. (**A**). Proto-arrays containing ∼9500 recombinant proteins were probed with either recombinant SETD3, SETD6 (His-SETD3 and His-SETD6) or BSA (negative control) followed by incubation with anti-His antibody and 647 Alexa fluor-conjugated secondary antibodies (**B**) Representative results are shown for PAK4, FOXM1 and AURKA. (**C**) Venn diagram showing the overlap between the identified SETD3 and SETD6 interacting proteins. Statistical significance of the overlap is shown. (**D**) Shared protein for SETD3 and SETD6 were categorized by biological process (bar graph) using the DAVID functional annotation tool.

We identified 176 SETD6-interacting proteins and 166 SETD3-interacting proteins. Representative proteins are shown in Figure 2B. The fact that both PAK4 and FoxM1 were previously characterized as substrates of SETD6 and SETD3, respectively, confirmed the reliability of this approach [10, 16] (Figure 2B). Out of the total proteins that were identified, 52 were found to be common for SETD3 and SETD6 (Figure 2C and Supplementary Table 1). Gene Ontology (GO) analysis of the shared proteins revealed a significant enrichment (P-value <0.05) of proteins involved in transcription, mRNA processing, RNA splicing, insulin secretion, and mitotic division (Figure 2D).

### SETD3 and SETD6 Target common Genes and Biological Processes

Due to the significant enrichment of interacting partners involved in transcription in the GO analysis, in addition to previous data linking both enzymes to transcription regulation [6, 8, 9, 16, 17], we hypothesized that their functional redundancy is reflected in shared target genes. To address this, we have generated HeLa cells depleted of SETD3, SETD6 and both, using the CRISPR-Cas9 system (Figure 3A), followed by an RNA-seq experiment. The principal component analysis (PCA) shows that all samples are distinct from control cells (Figure 3B). Double KO cells were the most distant from control cells (approximately 60% variability in PC1). Surprisingly, they were also distant from any of the single KO cells. The fact that the double KO displayed distinct transcriptomic profiles from any of the single KO cells suggests a functional link between the two enzymes.

**Figure 3.**
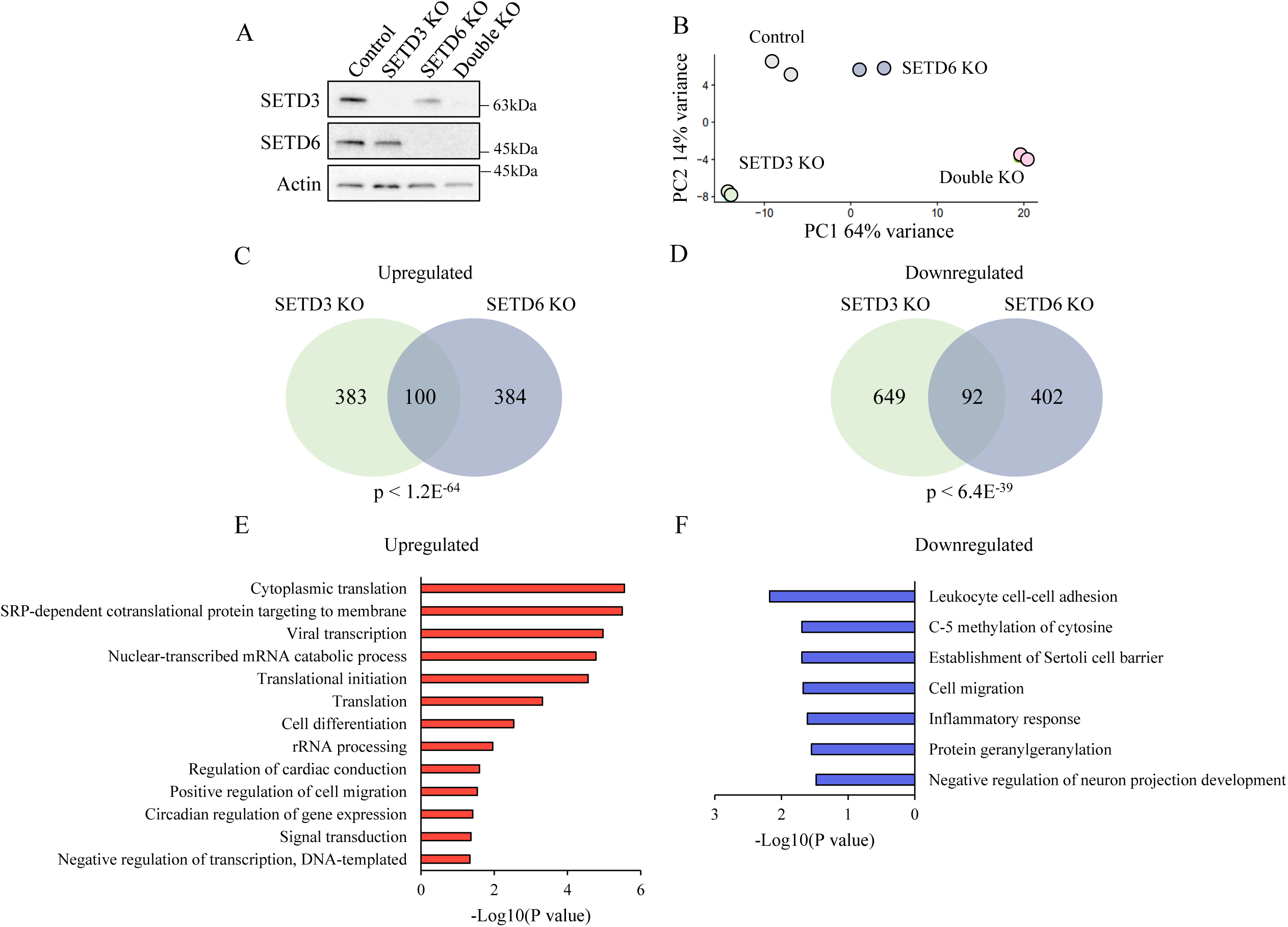
SETD3 and SETD6 share common target genes and biological processes. **(A)** Western Blot analysis using the indicated antibodies for HeLa control, SETD3 knock-out (SETD3 KO), SETD6 knock-out (SETD6 KO) and double knock-out (Double KO) cells that were generated using the CRISPR-Cas9 system (**B**) Principal component analysis (PCA) for cells shown A that were subjected for RNA-seq analysis. **(C+D)** Venn diagram showing the number of upregulated (C) or downregulated (D) genes (Adj. P-value <0.05), of each KO cell type compared to control cells. Statistical significance of the overlap is showing. **(E+F)** Shared upregulated (E) and downregulated (F) genes were categorized by biological processes using the DAVID annotation tool.

We first focused on the single KO cells to identify shared target genes. SETD6 depletion resulted in 484 upregulated and 494 downregulated genes. SETD3 KO cells had 483 upregulated and 741 downregulated genes (Adj P value <0.05). Consistent with our hypothesis, they shared 100 upregulated genes (Figure 3C), and 92 downregulated genes (Figure 3D). We further analyzed the shared genes for GO biological processes. In the upregulated genes (Figure 3E), we found significant enrichment of processes such as translation, viral transcription, cell differentiation and migration. In the down regulated genes (Figure 3F), we found an enrichment of processes such as cell-cell adhesion, cytosine methylation, inflammatory response and cell migration. Most of the identified processes were previously reported as SETD6 or SETD3-related [6, 11, 15, 35]. Taken together, our results indicate that SETD3 and SETD6 regulate these processes through common target genes.

### SETD3 and SETD6 Double Depletion Results in Increased Apoptosis

We next analyzed the double KO cells. We speculated that if SETD3 and SETD6 share common cellular functions, we might not reveal it in cells depleted of only one enzyme due to compensation by the other. Thus, we were especially interested in genes that changed in the double KO cells (Figure 4A). Indeed, we observed a dramatic change in the number of differentially expressed genes in comparison to the numbers obtained in the single KO cells (Figure 4B). GO analysis for the downregulated genes revealed enrichment of processes such as mRNA stability, amino-acid metabolism and NFκb signaling (Figure 4C). For the upregulated genes we found a significant enrichment of genes related to apoptosis and specifically DNA-damaged induced apoptosis (Figure 4D). A full list of the differentially expressed genes can be found in Supplementary Table 2.

**Figure 4.**
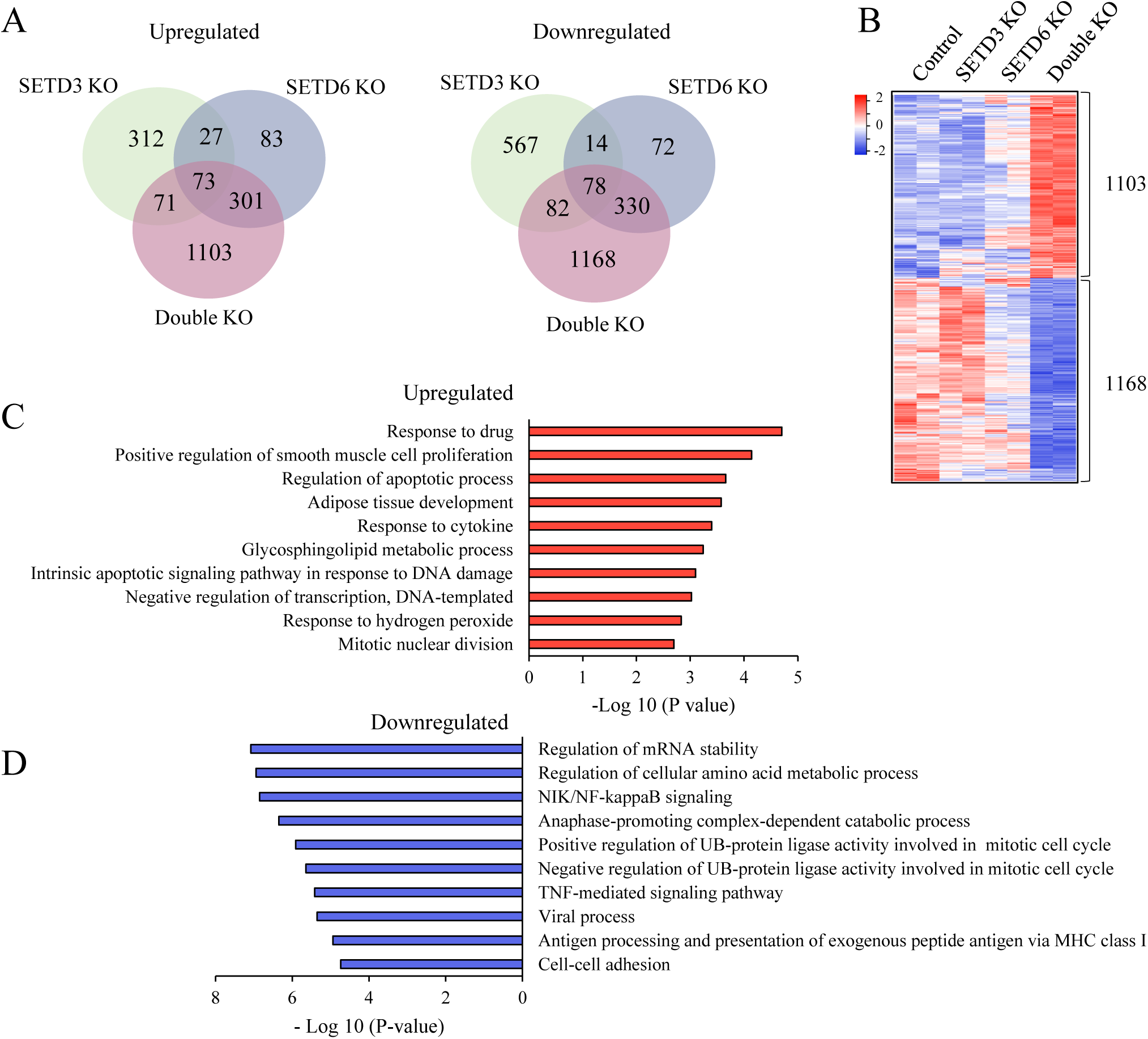
Double KO cells exhibit high level of apoptosis-related genes. **(A)** Venn diagram showing the number of upregulated and downregulated genes of the different KO cell types compared to control cells (Adj. P-value <0.05). (**B**) Heat map of 2271 genes which their expression significantly changed only in the double KO cells. Color bar represents high (Red) and low (Blue) expression levels. (**C+D**) Bar diagram showing biological processes enriched in the up-regulated (C) and down-regulated (D) genes as analyzed by the DAVID functional annotation tool.

As both SETD3 and SETD6 were shown before to be involved in the regulation of cell proliferation, cell cycle and apoptotic related cellular pathways [12, 17, 36-38], we speculated that the double KO cells will display substantial phenotypic alterations related to these processes. To address this hypothesis, we first successfully validated by qPCR the expression of several related genes that were found to be significantly upregulated in the double KO cells in the RNA-seq experiment (Figure 5A). We next tested if the increased expression of these genes in the double KO cells has a functional cellular phenotypic outcome. As shown in figure 5B, we did not detect significant differences in the proliferation rate between control and double KO cells. To study the involvement of SETD3 and SETD6 in cell cycle progression, the control and the double KO cells were synchronized by double thymidine block (G1/S) and the DNA content reflecting the different cell cycle stages, was measured by FACS (Figure 5C). The results revealed that the double KO cells progress slightly faster through the cell cycle compared to the control cells. At 8 h post thymidine release, ∼10% of the double KO cells enter G1, compared to 4% of the control cells. However, the effect was similar to previously reported data of SETD6 KO cells [12], suggesting that this observation is SETD6-dependant and not SETD3. Thus, we concluded that SETD3 is probably not involved in the regulation of cell-cycle in these cells. Finally, we tested whether the increased expression of apoptotic genes will lead to high apoptosis levels in the double KO cells. To this end, cells were treated with the DNA-damage inducer Doxorubicin for 24 hours and stained for Annexin-V and DAPI. Interestingly, regardless of the treatment, the basal apoptosis level in the double KO cells was slightly higher compared to the single KO cells. Similarly, the apoptotic levels following Doxorubicin induction were significantly higher in the double KO cells compared to the control and each of the single-KO cells (Figure 5D+E). Altogether, our results suggest a potential common function of SETD6 and SETD3 in apoptosis regulation.

**Figure 5.**
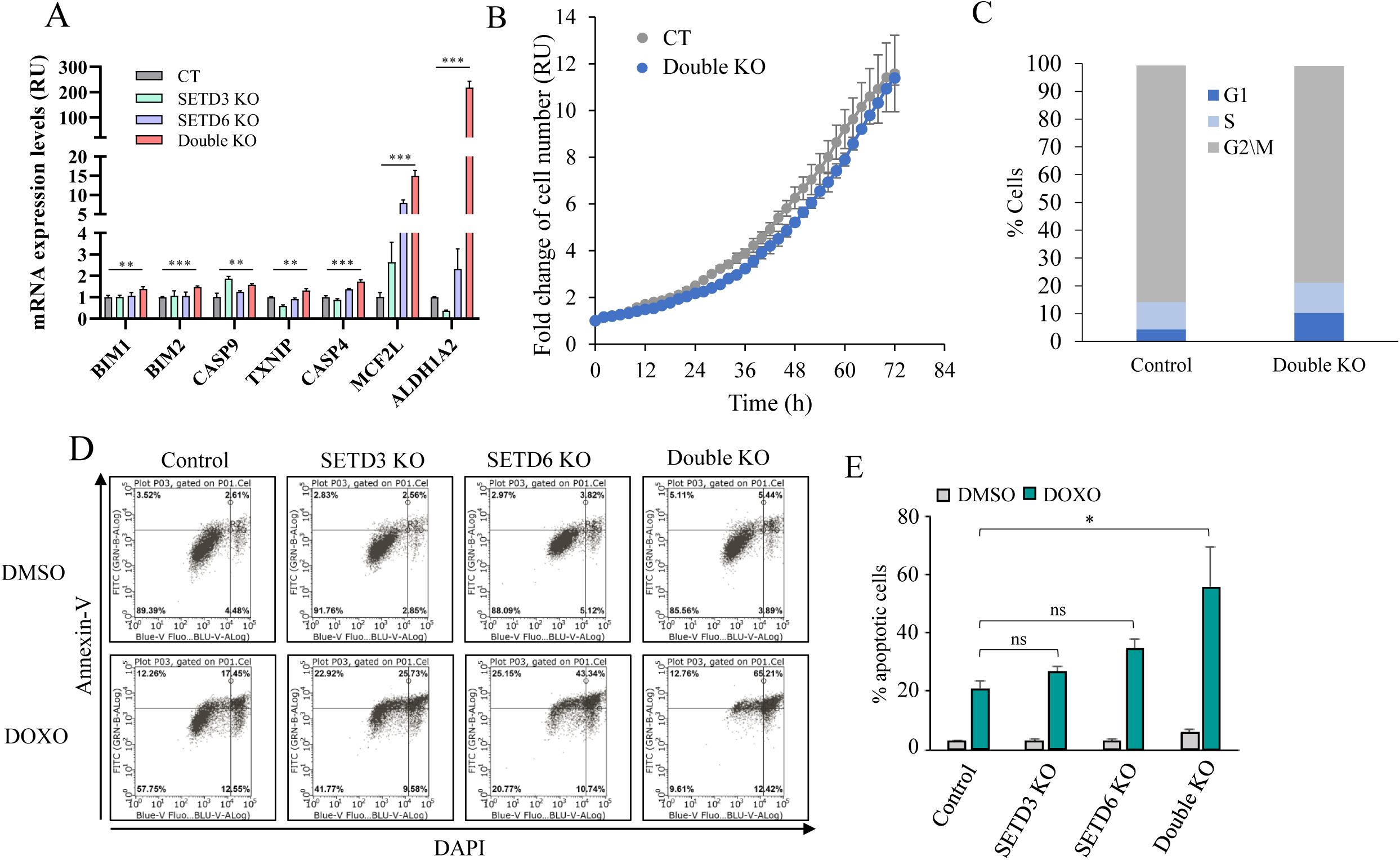
Cells depleted of both SETD6 and SETD3 show high apoptotic levels. (**A**) Validation by qPCR of representative genes (normalized to GAPDH), identified in the RNA-sequencing. Statistical analysis was performed for three experimental repeats using student’s t-test (**P < 0.01, ***P<0.001). (**B**) Cell proliferation assay of HeLa control and double KO cells. Cell number was monitored and calculated by LionheartTM FX Automated Microscope (4x) every 2 hours. Error bars are s.d. (**C**) Cell-cycle assay. HeLa control and double KO cells were synchronized using the double thymidine block method. Cells were harvested, fixed and stained with PI and then subjected to FACS analysis. The graph shows % of cells in G1, S and G2/M. (**D**) FACS analysis of HeLa control and KO cells treated with 2μM Doxorubicin (DOXO) or DMSO for 24h followed by staining with DAPI and Annexin V. (**E**) Quantification of % apoptotic cells (Annexin-V+ DAPI positive cells) from 3 independent FACS experiments. * p≤0.05

## Discussion

SETD6 and SETD3 are evolutionary relatives with a significant structure similarity. This finding raised the hypothesis that they share a similar role in cells. Our proteomic and genomic screenings revealed common interacting partners and target genes for SETD6 and SETD3. Moreover, cells depleted of both enzymes exhibited a dramatic change in the expression of genes related to several oncogenic-related cellular functions.

Our observation that 52 proteins may interact with both SETD6 and SETD3 strongly suggests that they have common substrates and raises the question on how they share cellular functions. One possibility is that they methylate the same residue. Another option is that they methylate different residues of the same substrate. Recent studies claim that SETD3 is a histidine methyltransferase, with no lysine methylation activity [22, 23, 34, 39]. Thus, it is more likely that they methylate different residues. However, an intriguing question that remains open, is to understand whether and how these two methylation events on two different residues work in concert or separately.

Our RNA-seq data revealed many genes that did not significantly change in the single KO cells but changed dramatically in the double KO cells. By focusing on these genes, we could identify cases in which one of the enzymes compensated for the loss of the other. GO analysis of these genes revealed enrichment of several processes, among them the apoptotic pathway. Our results indicate that the double KO cells led to high apoptotic levels both on both, basal and under DNA damage conditions. Our data further indicates that SETD3 and SETD6 negatively regulate apoptosis through the inhibition of apoptosis-related genes. However, we still do not understand the molecular mechanism underlying these observations. The fact that SETD6 and SETD3 share interacting proteins implies for a common substrate. However, there is still a possibility that they methylate different substrates in the same pathway, leading to similar consequences. SETD6 has previously been shown to increase cell survival in bladder cancer and kidney cells [37, 38], which supports our findings. However, SETD3 has been found to promote apoptosis in colon cancer cells in a p53 dependent manner [17]. This observation can be explained by the fact that HeLa cells do not have functional p53 [40, 41], and this mechanism might be irrelevant in these cells.

Evading apoptosis is one of the main hallmarks of cancer, and apoptosis induction is a major target for therapeutic approaches [41]. Our results suggest that SETD6 and SETD3 inhibit apoptosis through gene regulation. Thus, inhibition of both might be relevant as a cancer-treatment strategy. Given the high structural similarity between SETD6 and SETD3, it might be possible to design a single inhibitor for both. In recent years, lysine methyltransferases have become attractive candidates for drug design. For example, several G9a inhibitors were designed and displayed an anti-tumor effect in mouse models [42]. EZH2 inhibitor Tazemetostat is in phase II trials against relapsed or refractory non-Hodgkin lymphoma [43]. DOT1L inhibitor EPZ-5676 is also undergoing phase I/II against Relapsed/Refractory Leukemia [44]. Recently, a β-actin based-peptide was designed as an inhibitor for SETD3 [45]. However, there is still no available data regarding its effect on cells.

The enzymatic activity and substrate recognition of SETD6 and SETD3 have been studied previously. However, to the best of our knowledge this is the first report of the functional crosstalk between the two enzymes. Our results strongly suggest a shared function of SETD6 and SETD3 in regulating DNA-damage-induced apoptosis and set the base for future studies combining protein methylation and cancer therapy.

## Acknowledgments

We thank the Levy lab for technical assistance and helpful discussions. This work was supported by grants to DL from The Israel Science Foundation (285/14 and 262/18), Israel Cancer Association) ICA(and from the Israel Cancer Research Fund (ICRF).

## Author Contribution

LE, EAS, OC MF and DL designed and performed the experiments. GS and NS performed the bioinformatic analysis. LE and DL wrote the paper. All authors read and approved the final manuscript.

## Conflict of Interest

The authors declare that they have no conflict of interest.

## Notes

### Competing Interest Statement

The authors have declared no competing interest.

## References

[1] M. Albert, K. Helin, Histone methyltransferases in cancer, Seminars in cell & developmental biology, 21 (2010) 209–220.

[2] C.H. Arrowsmith, C. Bountra, P.V. Fish, K. Lee, M. Schapira, Epigenetic protein families: a new frontier for drug discovery, Nature reviews. Drug discovery, 11 (2012) 384–400.

[3] P.O. Falnes, M.E. Jakobsson, E. Davydova, A. Ho, J. Malecki, Protein lysine methylation by seven-beta-strand methyltransferases, Biochem J, 473 (2016) 1995–2009.

[4] S.G. Clarke, 16 Inhibition of mammalian protein methyltransferases by 5’-methylthioadenosine (MTA): A mechanism of action of dietary same?, Enzymes, 24 (2006) 467–493.

[5] R. Hamamoto, V. Saloura, Y. Nakamura, Critical roles of non-histone protein lysine methylation in human tumorigenesis, Nature reviews. Cancer, 15 (2015) 110–124.

[6] D. Levy, A.J. Kuo, Y. Chang, U. Schaefer, C. Kitson, P. Cheung, A. Espejo, B.M. Zee, C.L. Liu, S. Tangsombatvisit, R.I. Tennen, A.Y. Kuo, S. Tanjing, R. Cheung, K.F. Chua, P.J. Utz, X. Shi, R.K. Prinjha, K. Lee, B.A. Garcia, M.T. Bedford, A. Tarakhovsky, X. Cheng, O. Gozani, Lysine methylation of the NF-kappaB subunit RelA by SETD6 couples activity of the histone methyltransferase GLP at chromatin to tonic repression of NF-kappaB signaling, Nature immunology, 12 (2011) 29–36.

[7] D.J. O’Neill, S.C. Williamson, D. Alkharaif, I.C. Monteiro, M. Goudreault, L. Gaughan, C.N. Robson, A.C. Gingras, O. Binda, SETD6 controls the expression of estrogen-responsive genes and proliferation of breast carcinoma cells, Epigenetics, 9 (2014) 942–950.

[8] O. Binda, A. Sevilla, G. LeRoy, I.R. Lemischka, B.A. Garcia, S. Richard, SETD6 monomethylates H2AZ on lysine 7 and is required for the maintenance of embryonic stem cell self-renewal, Epigenetics, 8 (2013) 177–183.

[9] A. Chen, M. Feldman, Z. Vershinin, D. Levy, SETD6 is a negative regulator of oxidative stress response, Biochimica et biophysica acta, 1859 (2016) 420–427.

[10] Z. Vershinin, M. Feldman, A. Chen, D. Levy, PAK4 Methylation by SETD6 Promotes the Activation of the Wnt/beta-Catenin Pathway, The Journal of biological chemistry, 291 (2016) 6786–6795.

[11] Z. Vershinin, M. Feldman, T. Werner, L.E. Weil, M. Kublanovsky, E. Abaev-Schneiderman, M. Sklarz, E.Y.N. Lam, K. Alasad, S. Picaud, B. Rotblat, R.A. McAdam, V. Chalifa-Caspi, M. Bantscheff, T. Chapman, H.D. Lewis, P. Filippakopoulos, M.A. Dawson, P. Grandi, R.K. Prinjha, D. Levy, BRD4 methylation by the methyltransferase SETD6 regulates selective transcription to control mRNA translation, Sci Adv, 7 (2021).

[12] M. Feldman, Z. Vershinin, I. Goliand, N. Elia, D. Levy, The methyltransferase SETD6 regulates Mitotic progression through PLK1 methylation, Proc Natl Acad Sci U S A, 116 (2019) 1235–1240.

[13] V.M. Richon, D. Johnston, C.J. Sneeringer, L. Jin, C.R. Majer, K. Elliston, L.F. Jerva, M.P. Scott, R.A. Copeland, Chemogenetic analysis of human protein methyltransferases, Chem Biol Drug Des, 78 (2011) 199–210.

[14] D.W. Kim, K.B. Kim, J.Y. Kim, S.B. Seo, Characterization of a novel histone H3K36 methyltransferase setd3 in zebrafish, Biosci Biotechnol Biochem, 75 (2011) 289–294.

[15] G.H. Eom, K.B. Kim, J.H. Kim, J.Y. Kim, J.R. Kim, H.J. Kee, D.W. Kim, N. Choe, H.J. Park, H.J. Son, S.Y. Choi, H. Kook, S.B. Seo, Histone methyltransferase SETD3 regulates muscle differentiation, J Biol Chem, 286 (2011) 34733–34742.

[16] O. Cohn, M. Feldman, L. Weil, M. Kublanovsky, D. Levy, Chromatin associated SETD3 negatively regulates VEGF expression, Scientific reports, 6 (2016) 37115.

[17] E. Abaev-Schneiderman, L. Admoni-Elisha, D. Levy, SETD3 is a positive regulator of DNA-damage-induced apoptosis, Cell Death Dis, 10 (2019) 74.

[18] L. Xu, P. Wang, X. Feng, J. Tang, L. Li, X. Zheng, J. Zhang, Y. Hu, T. Lan, K. Yuan, Y. Zhang, S. Ren, X. Hao, M. Zhang, M. Xu, SETD3 is regulated by a couple of microRNAs and plays opposing roles in proliferation and metastasis of hepatocellular carcinoma, Clin Sci (Lond), 133 (2019) 2085–2105.

[19] A.S. Pires-Luis, M. Vieira-Coimbra, F.Q. Vieira, P. Costa-Pinheiro, R. Silva-Santos, P.C. Dias, L. Antunes, F. Lobo, J. Oliveira, C.S. Goncalves, B.M. Costa, R. Henrique, C. Jeronimo, Expression of histone methyltransferases as novel biomarkers for renal cell tumor diagnosis and prognostication, Epigenetics, 10 (2015) 1033–1043.

[20] N. Hassan, N. Rutsch, B. Gyorffy, N.A. Espinoza-Sanchez, M. Gotte, SETD3 acts as a prognostic marker in breast cancer patients and modulates the viability and invasion of breast cancer cells, Sci Rep, 10 (2020) 2262.

[21] H. Engqvist, T.Z. Parris, A. Kovacs, E.W. Ronnerman, K. Sundfeldt, P. Karlsson, K. Helou, Validation of Novel Prognostic Biomarkers for Early-Stage Clear-Cell, Endometrioid and Mucinous Ovarian Carcinomas Using Immunohistochemistry, Front Oncol, 10 (2020) 162.

[22] A.W. Wilkinson, J. Diep, S. Dai, S. Liu, Y.S. Ooi, D. Song, T.M. Li, J.R. Horton, X. Zhang, C. Liu, D.V. Trivedi, K.M. Ruppel, J.G. Vilches-Moure, K.M. Casey, J. Mak, T. Cowan, J.E. Elias, C.M. Nagamine, J.A. Spudich, X. Cheng, J.E. Carette, O. Gozani, SETD3 is an actin histidine methyltransferase that prevents primary dystocia, Nature, 565 (2019) 372–376.

[23] S. Kwiatkowski, A.K. Seliga, D. Vertommen, M. Terreri, T. Ishikawa, I. Grabowska, M. Tiebe, A.A. Teleman, A.K. Jagielski, M. Veiga-da-Cunha, J. Drozak, SETD3 protein is the actin-specific histidine N-methyltransferase, Elife, 7 (2018).

[24] A.H. Peters, D. O’Carroll, H. Scherthan, K. Mechtler, S. Sauer, C. Schofer, K. Weipoltshammer, M. Pagani, M. Lachner, A. Kohlmaier, S. Opravil, M. Doyle, M. Sibilia, T. Jenuwein, Loss of the Suv39h histone methyltransferases impairs mammalian heterochromatin and genome stability, Cell, 107 (2001) 323–337.

[25] M. Tachibana, J. Ueda, M. Fukuda, N. Takeda, T. Ohta, H. Iwanari, T. Sakihama, T. Kodama, T. Hamakubo, Y. Shinkai, Histone methyltransferases G9a and GLP form heteromeric complexes and are both crucial for methylation of euchromatin at H3-K9, Genes Dev, 19 (2005) 815–826.

[26] P.A. Link, O. Gangisetty, S.R. James, A. Woloszynska-Read, M. Tachibana, Y. Shinkai, A.R. Karpf, Distinct roles for histone methyltransferases G9a and GLP in cancer germ-line antigen gene regulation in human cancer cells and murine embryonic stem cells, Mol Cancer Res, 7 (2009) 851–862.

[27] N.T. Crump, T.A. Milne, Why are so many MLL lysine methyltransferases required for normal mammalian development?, Cell Mol Life Sci, 76 (2019) 2885–2898.

[28] D. Levy, C.L. Liu, Z. Yang, A.M. Newman, A.A. Alizadeh, P.J. Utz, O. Gozani, A proteomic approach for the identification of novel lysine methyltransferase substrates, Epigenetics Chromatin, 4 (2011) 19.

[29] M.Y. Sklarz, L. Levin, M. Gordon, V. Chalifa-Caspi, NeatSeq-Flow: A Friendly High-Throughput Sequencing Workflow Platform for Local Use by Programmers and Non-Programmers Alike, Eur J Hum Genet, 27 (2019) 1644–1645.

[30] A. Dobin, C.A. Davis, F. Schlesinger, J. Drenkow, C. Zaleski, S. Jha, P. Batut, M. Chaisson, T.R. Gingeras, STAR: ultrafast universal RNA-seq aligner, Bioinformatics, 29 (2013) 15–21.

[31] B. Li, C.N. Dewey, RSEM: accurate transcript quantification from RNA-Seq data with or without a reference genome, BMC Bioinformatics, 12 (2011) 323.

[32] P. Ewels, M. Magnusson, S. Lundin, M. Kaller, MultiQC: summarize analysis results for multiple tools and samples in a single report, Bioinformatics, 32 (2016) 3047–3048.

[33] M.I. Love, W. Huber, S. Anders, Moderated estimation of fold change and dispersion for RNA-seq data with DESeq2, Genome Biol, 15 (2014) 550.

[34] S. Dai, J.R. Horton, C.B. Woodcock, A.W. Wilkinson, X. Zhang, O. Gozani, X. Cheng, Structural basis for the target specificity of actin histidine methyltransferase SETD3, Nat Commun, 10 (2019) 3541.

[35] Z. Vershinin, M. Feldman, D. Levy, PAK4 methylation by the methyltransferase SETD6 attenuates cell adhesion, Sci Rep, 10 (2020) 17068.

[36] X. Cheng, Y. Hao, W. Shu, M. Zhao, C. Zhao, Y. Wu, X. Peng, P. Yao, D. Xiao, G. Qing, Z. Pan, L. Yin, D. Hu, H.N. Du, Cell cycle-dependent degradation of the methyltransferase SETD3 attenuates cell proliferation and liver tumorigenesis, J Biol Chem, 292 (2017) 9022–9033.

[37] N. Mukherjee, E. Cardenas, R. Bedolla, R. Ghosh, SETD6 regulates NF-kappaB signaling in urothelial cell survival: Implications for bladder cancer, Oncotarget, 8 (2017) 15114–15125.

[38] X. Wang, Q. Liu, D. Kong, Z. Long, Y. Guo, S. Wang, R. Liu, C. Hai, Down-regulation of SETD6 protects podocyte against high glucose and palmitic acid-induced apoptosis, and mitochondrial dysfunction via activating Nrf2-Keap1 signaling pathway in diabetic nephropathy, J Mol Histol, 51 (2020) 549–558.

[39] Q. Guo, S. Liao, S. Kwiatkowski, W. Tomaka, H. Yu, G. Wu, X. Tu, J. Min, J. Drozak, C. Xu, Structural insights into SETD3-mediated histidine methylation on beta-actin, Elife, 8 (2019).

[40] G. Matlashewski, L. Banks, D. Pim, L. Crawford, Analysis of human p53 proteins and mRNA levels in normal and transformed cells, Eur J Biochem, 154 (1986) 665–672.

[41] B.A. Carneiro, W.S. El-Deiry, Targeting apoptosis in cancer therapy, Nat Rev Clin Oncol, 17 (2020) 395–417.

[42] H. Cao, L. Li, D. Yang, L. Zeng, X. Yewei, B. Yu, G. Liao, J. Chen, Recent progress in histone methyltransferase (G9a) inhibitors as anticancer agents, Eur J Med Chem, 179 (2019) 537–546.

[43] N. Gulati, W. Beguelin, L. Giulino-Roth, Enhancer of zeste homolog 2 (EZH2) inhibitors, Leuk Lymphoma, 59 (2018) 1574–1585.

[44] C.R. Klaus, D. Iwanowicz, D. Johnston, C.A. Campbell, J.J. Smith, M.P. Moyer, R.A. Copeland, E.J. Olhava, M.P. Scott, R.M. Pollock, S.R. Daigle, A. Raimondi, DOT1L inhibitor EPZ-5676 displays synergistic antiproliferative activity in combination with standard of care drugs and hypomethylating agents in MLL-rearranged leukemia cells, J Pharmacol Exp Ther, 350 (2014) 646–656.

[45] J.C.J. Hintzen, L. Moesgaard, S. Kwiatkowski, J. Drozak, J. Kongsted, J. Mecinovic, beta-Actin Peptide-Based Inhibitors of Histidine Methyltransferase SETD3, ChemMedChem, 16 (2021) 2695–2702.

